# aRgus: multilevel visualization of non-synonymous single nucleotide variants & advanced pathogenicity score modeling for genetic vulnerability assessment

**DOI:** 10.1101/2022.10.20.513018

**Authors:** Julian Schröter, Tal Dattner, Jennifer Hüllein, Alejandra Jayme, Vincent Heuveline, Georg F. Hoffmann, Stefan Kölker, Dominic Lenz, Thomas Opladen, Bernt Popp, Christian P. Schaaf, Christian Staufner, Steffen Syrbe, Sebastian Uhrig, Daniel Hübschmann, Heiko Brennenstuhl

**Affiliations:** Division of Pediatric Epileptology, Center for Pediatrics and Adolescent Medicine, University Hospital Heidelberg, Im Neuenheimer Feld 430, D-69120 Heidelberg, Germany; Division of Neuropediatrics and Metabolic Medicine, Center for Pediatrics and Adolescent Medicine, University Hospital Heidelberg, Im Neuenheimer Feld 430, D-69120 Heidelberg, Germany; Computational Oncology, Molecular Precision Oncology Program, National Center for Tumor Diseases (NCT), German Cancer Research Center (DKFZ), Im Neuenheimer Feld 460, D-69120 Heidelberg, Germany; Engineering Mathematics and Computing Lab (EMCL), Interdisciplinary Center for Scientific Computing (IWR), University of Heidelberg, Im Neuenheimer Feld 205, D-69120 Heidelberg, Germany; Institute of Human Genetics, University Medical Center Leipzig, Philipp-Rosenthal-Str. 55 (Haus W), D-04103 Leipzig, Germany; Institute of Human Genetics, University Hospital Heidelberg, Im Neuenheimer Feld 440, D-69120 Heidelberg, Germany; German Cancer Consortium (DKTK), Im Neuenheimer Feld 280, D-69120 Heidelberg, Germany; Heidelberg Institute for Stem Cell Technology and Experimental Medicine (HI-STEM), Im Neuenheimer Feld 280, D-69120 Heidelberg, Germany

**Keywords:** Pathogenicity scores, variant effect prediction, variant assessment, computational genetics

## Abstract

The widespread use of high-throughput sequencing techniques is leading to a rapidly increasing number of disease-associated variants of unknown significance and candidate genes. Integration of knowledge concerning their genetic, protein as well as functional and conservational aspects is necessary for an exhaustive assessment of their relevance and for prioritization of further clinical and functional studies investigating their role in human disease. In order to collect the necessary information, a multitude of different databases has to be accessed and data extraction from the original sources commonly is not user-friendly and requires advanced bioinformatics skills. This leads to a decreased data accessibility for a relevant number of potential users such as clinicians, geneticist, and clinical researchers. Here, we present aRgus (https://argus.urz.uni-heidelberg.de/), a standalone webtool for simple extraction and intuitive visualization of multi-layered gene, protein, variant, and variant effect prediction data. aRgus provides interactive exploitation of these data within seconds for any known gene of the human genome. In contrast to existing online platforms for compilation of variant data, aRgus complements visualization of chromosomal exon-intron structure and protein domain annotation with ClinVar and gnomAD variant distributions as well as position-specific variant effect prediction score modeling. aRgus thereby enables timely assessment of protein regions vulnerable to variation with single amino acid resolution and provides numerous applications in variant and protein domain interpretation as well as in the design of *in vitro* experiments.

## 1. Introduction

In recent years, high-throughput sequencing methods have led to a tremendous increase in the extent of genetic and variant data related to human disease (1, 2). Upon identification of disease-associated genetic variants of unknown significance or in novel candidate genes, an investigator may need to integrate of multi-layered information concerning exon-intron structure, protein domain annotation, mutational constraint, as well as known variants present in patients and healthy individuals including their allele frequency. Additionally, the potential biological impact of variants on protein structure and function can be predicted using *in silico* pathogenicity scores that assign a numerical value to each amino acid substitution. This is particularly helpful for estimation of damaging variant effects when no functional *in vitro* data is available. This information has to be taken into consideration for variant interpretation according to the ACMG guidelines (3). Although the majority of the above-mentioned data are publicly accessible, they are only available in abstract, tabular form, stored in a multitude of different databases that have to be accessed individually, and their extraction, formatting, and analysis often require extensive bioinformatic capabilities. User-friendly platforms have previously been developed in order to facilitate access to genetic data from several resources but lack detailed integration and visualization of different pathogenicity scoring models (4–7).

Therefore, we developed aRgus (https://argus.urz.uni-heidelberg.de/) as a standalone webtool for user-friendly and intuitive compilation and visualization of complex data on genetic variants and *in silico* pathogenicity scores from the extensive databases Ensembl, Simple ClinVar, the Universal Protein Resource (UniProt), the Genome Aggregation Database (gnomAD), and dbNSFP (4, 5, 7–9). The Ensembl database contains comprehensive genomic information including chromosomal gene and transcript localization (4). Simple ClinVar is an interactive webtool using a custom algorithm to retrieve simplified summary statistics on variant and phenotype information from ClinVar, the largest archive of genetic variants associated with human disease (5, 10). UniProt represents the largest database for protein sequence and domain annotation data (7). The gnomAD database contains variant data from nearly 150,000 healthy individuals identified in exome and genome sequencing studies (8). The dbNSFP database represents a rich resource containing values of numerous *in silico* pathogenicity scores precalculated for all biologically possible non-synonymous single-nucleotide variants (nsSNVs) and related information, such as their gnomAD allele frequencies, that can be used for variant annotation (9). dbNSFP is implemented in several annotation tools such as ANNOVAR, VarSome, the UCSC Genome Browser, and the Ensembl Variant Effect Predictor and also offers an own application but can only be used for single queries or short lists of SNVs (6, 11–13).

In contrast, aRgus provides the synopsis of both variant and pathogenicity score data using an intuitive graphical user interface. aRgus allows display of exon-intron structure and protein domain annotation together with ClinVar and gnomAD variant distributions, a vivid visualization of pathogenicity score values and their statistical comparison in different variant groups, as well as an interactive table comprising ClinVar- and dbNSFP-derived variants. The use of aRgus enables identification of protein regions susceptible to missense variation up to single amino acid (AA) resolution and represents a powerful tool for enhanced inference-based variant interpretation.

## 2. Methods

### 2.1. Implementation

aRgus is implemented as a standalone application using the RStudio shiny framework (https://shiny.rstudio.com/) that allows translation of remote user operations into HTML code. Chromosomal coordinates and the UniProt ID of the transcript are retrieved through Ensembl (14) using the R package *AnnotationHub*. In order to achieve user-friendliness and to maximize the quality of data retrieval, the canonical transcript is automatically determined via query of the MANE transcript (15) or the highest quality APPRIS isoform (16). ClinVar variant and phenotype annotation is retrieved using a monthly updated dataset generated via the Simple ClinVar filter (5). Domain and region annotations of the corresponding protein are directly retrieved from UniProt using the R package *drawProteins* (7, 17). We use a tabix-indexed dbNSFP (v.4.3a) file to access up to 43 *in silico* pathogenicity scores for all possible nsSNVs and their gnomAD (exomes v.2.1, genomes v.3.0) allele counts (9). All databases are updated in regular intervals according to their respective release cycle. All visualizations are realized using the R library ggplot2 v3.2.1 (18). Each plot (.svg/.png) and table (.csv/.xlsx) can be exported separately for offline data processing. The aRgus web server is compatible with all common web browser applications including versions for mobile devices. The source code is available at https://github.com/huellejn/argus. The application can be deployed locally using a Docker image.

### 2.2. Visualization of tabular pathogenicity score data

Theoretically, a gene transcript can mutate at any base position into three alternate bases leading to nsSNVs on the gene level as well as amino acid substitutions or truncations on the protein level, depending on the position within the base triplet. The damaging effect on protein function can be predicted *in silico* by an individual value of different pathogenicity scores assigned to each amino acid substitution (Fig. S1). Thus, all biologically possible nsSNVs can be simulated and result in several datapoints per amino acid position. In order to visualize these data intuitively and vividly, a dual approach was conducted: First, the *geom_smooth()* function of the R package *ggplot2* was used to generate a polynomial regression of smoothed conditional means displayed by an approximation curve with 95% confidence interval. Local Polynomial Regression Fitting (*loess*, formula = y ~ x) and a generalized additive model (*GAM*, formula = y ~ s(x, bs = “cs”)) are used for < and ≥ 1,000 datapoints, respectively. Second, the arithmetic means of multiple pathogenicity score values at one amino acid position were calculated and visualized as a heat-strip color-coded by the predicted degree of the damaging effect on protein function (Fig. S1).

### 2.3. Statistics

All pathogenicity scores can be subjected to t-test comparisons between four pre-defined groups: 1.) variants stored in ClinVar and classified as pathogenic/likely pathogenic (*ClinVar_pathogenic*), 2.) variants stored in ClinVar and classified as benign/likely benign (*ClinVar_benign*), 3.) variants stored in gnomAD (*gnomAD*), and 4.) all biologically possible variants stored in dbNSFP (*InSilico*). Score value distributions within these groups are displayed as violin plots with integrated quartiles. The level of significance is shown as asterisks as follows: * = p < 0.05, ** = p < 0.01, *** = p < 0.001.

## 3. Results

### 3.1. Main user interface

aRgus provides intuitive use and accessibility. It can be accessed via all common browsers and operating systems including mobile devices. On the aRgus main page, the user can enter a gene of interest via its HGNC symbol (Fig. 1) The tool subsequently provides the MANE- and APPRIS-curated canonical transcripts. The user can then choose from a panel of six plots that can be displayed in a modular way in order to allow an individual compilation: 1.) Unspliced transcript plot; 2.) protein plot; 3.) mutational constraint plots of disease-associated and putatively benign ClinVar as well as 4.) tolerated gnomAD variants; 5.) a combined pathogenicity score model including a polynomial fit and heat-strip with position-coded annotation of score mean values; and 6.) a statistical comparison of score values of different variant groups. Additionally, two interactive tables are available including a tab for all ClinVar variants (*ClinVar*) and all biologically possible nsSNVs together with corresponding score values derived from the dbNSFP database (*In silico scores*), respectively. Plots can be exported in two file formats (.png/.svg) with user-specified aspect ratios. Tables can be exported as .csv or .xlsx files for individual data storage and further offline data manipulation.

**Fig. 1:**
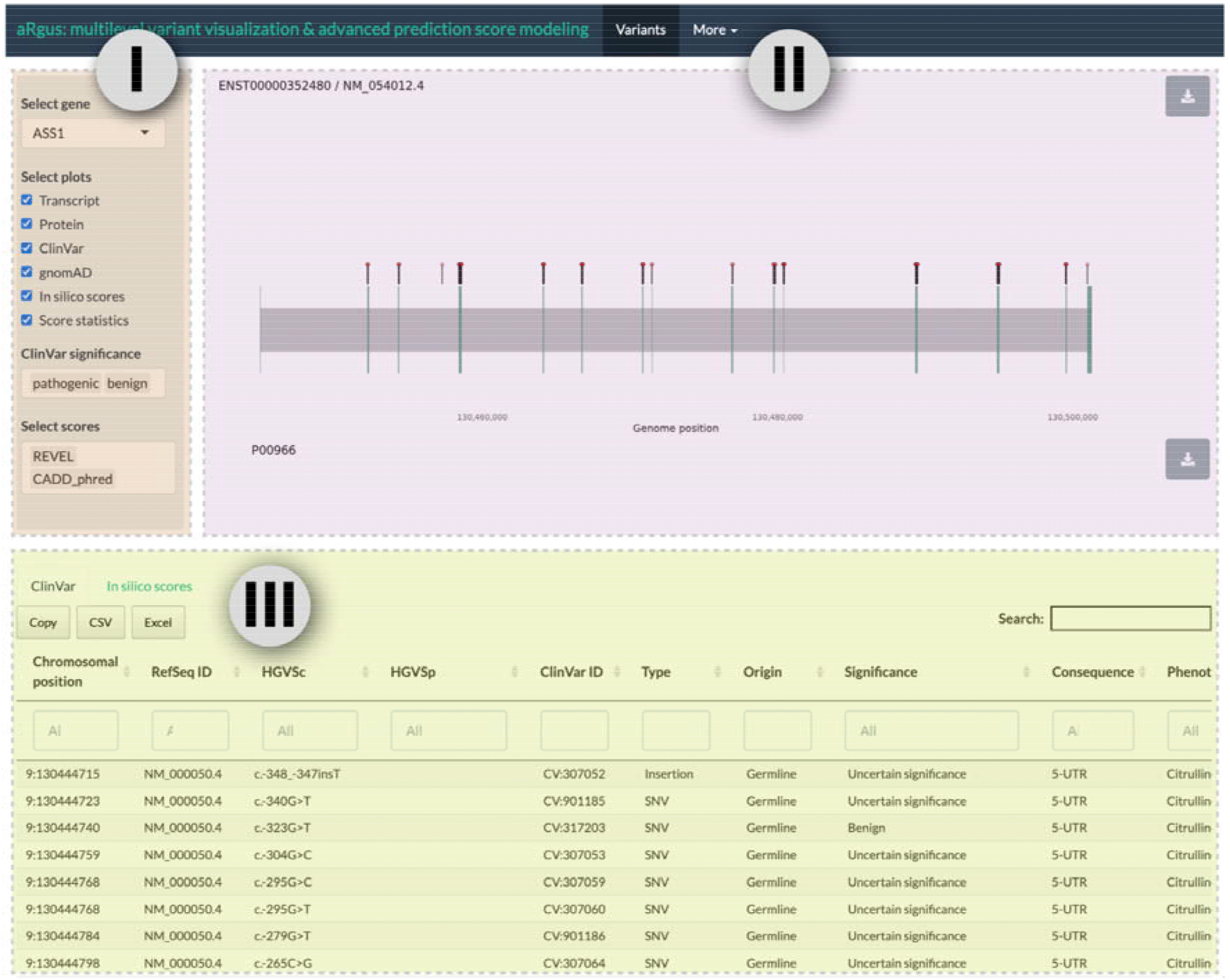
aRgus user interface. I) Interactive input mask with control elements, II) dynamic results area, III) tables from which variants can be selected for display with label.

### 3.2. Applications

#### 3.2.1. Unspliced transcript plot

The unspliced transcript plot (UTP) displays the gene’s scaled exon-intron structure from left to right starting with the first exon for improved readability regardless of the genomic localization on the forward or reverse strand. By default, pathogenic and likely pathogenic (P/LP) ClinVar variants are shown as lollipops which allows convenient visualization of intronic variants. In order to display the variant description, ClinVar and simulated dbNSFP variants can be manually selected in the respective tables. Figure 2A shows the UTP for the gene *ASS1*, encoding the enzyme argininosuccinate synthase (ASS), with selected P/LP ClinVar variants (red) and variants from the *In silico scores* table (gray), containing the dbNSFP-derived variants.

**Fig. 2:**
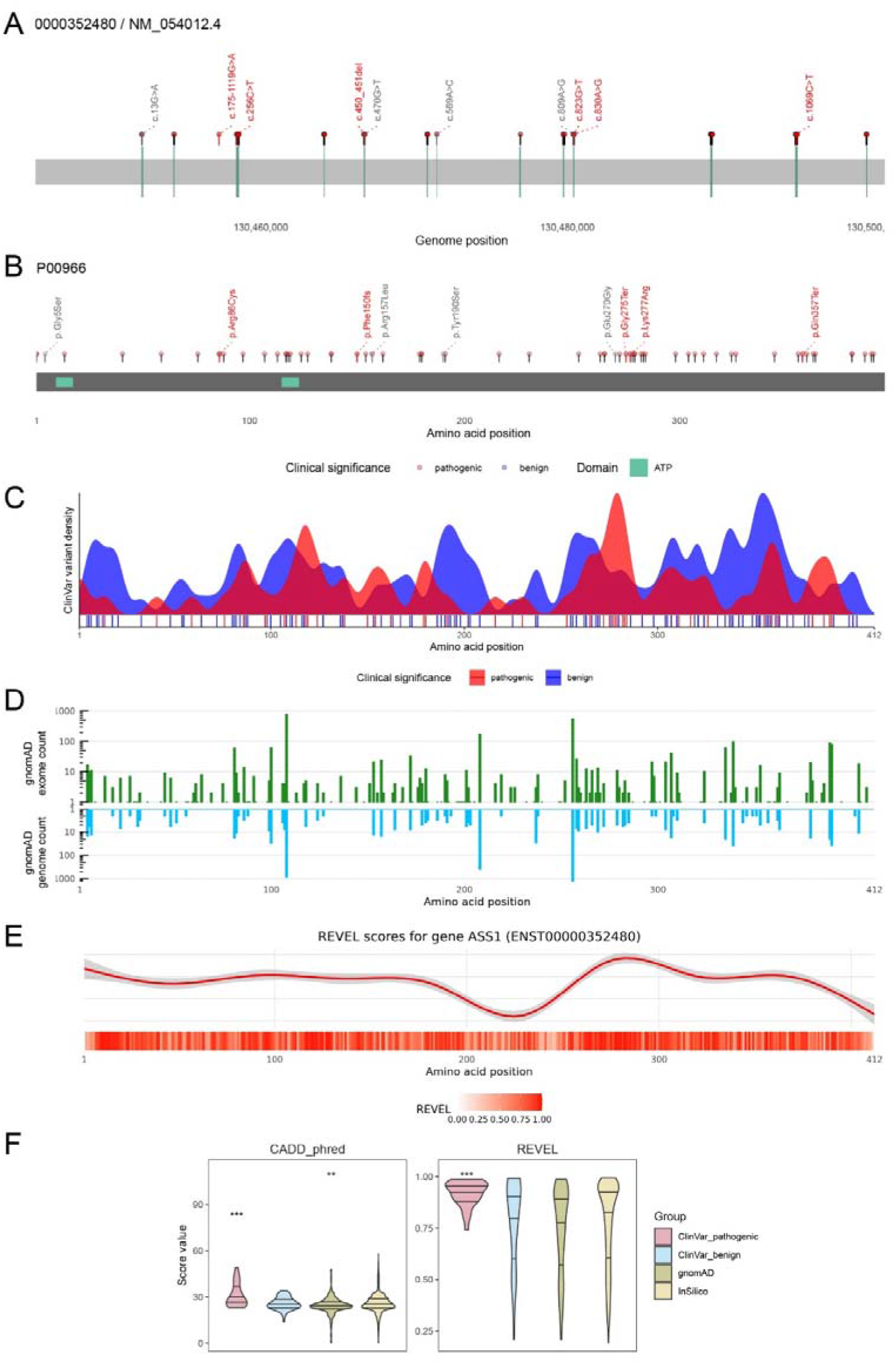
aRgus plots. A) UTP of the gene *ASS1*. Labels show P/LP variants (red) and selected variants from the *in silico* tab (gray). B) Protein plot with AA exchanges corresponding to variants shown in A). C) Density plot of P/LP (red) and benign/likely benign (blue) Simple ClinVar variants. D) Logarithmic histogram of gnomAD exomes (green) and genomes (blue) variant allele frequencies. E) Polynomial regression of REVEL score (top) and heat-strip of mean score values (bottom). F) t-test group comparisons shown as violin plots with quartiles, * (*p*-value < 0.05), ** (*p*-value < 0.01), and *** (*p*-value < 0.001).

#### 3.2.2. Protein plot

The primary structure of the resulting protein is visualized by the protein plot showing a linearized representation together with annotated domains retrieved from UniProt. As in the UTP, variants can be manually selected from the provided tables. Thereby, distribution of known and novel variants and their relation to protein domains/regions can easily be assessed. This versatile visualization provides useful insights for assessment of the pathophysiological relevance of potentially functionally relevant domains, given a gene scarcely associated with pathogenic variants. Figure 2B shows respective amino acid changes and protein domains of ASS.

#### 3.2.3. ClinVar and gnomAD mutational constraint plots

Distributions of ClinVar and gnomAD variants with respect to their protein position and allele frequency are visualized by density and bar plots, respectively, facilitating assessment of a protein’s mutational constraint. This includes sections of mutational hotspots, recurrent pathogenic and benign variants as well as the position-specific degree of tolerance towards missense variation. For more precise localization, ClinVar variants are additionally shown as vertical lines underneath the density curves (Fig. 2C). gnomAD variants are displayed in two separate logarithmic bar plots depending on their origin from the exomes (green) or genomes (blue) dataset (Fig. 2D). For ASS, ClinVar density curves reveal an accumulation of pathogenic variants in the region of AA 260-280 whereas gnomAD variants from both exomes and genomes show low population allele frequencies or are completely absent from the dataset (Figure 2E).

#### 3.2.4. *In silico* pathogenicity score model

Pre-calculated pathogenicity score values of all biologically possible nsSNVs are retrieved from the dbNSFP database. In order to improve data accessibility, the resulting multiple data points per protein position are simplified and visualized using a polynomial regression model combined with a heat-strip scaled to the linear protein representation. Depending on the user’s research question, the desired pathogenicity scoring model can immediately be selected from a list of up to 43 different scores. Plots for three different scores can be displayed simultaneously. This enables assessment of the predicted, position-specific impact of amino acid substitutions within the context of known protein domains and facilitates detection of regions of increased or decreased susceptibility to missense variation. Thereby, the functional impact of novel variants can be estimated and investigation of unknown sections of predicted damaging variant effects can be addressed in order to formulate future research hypotheses.

In our practical example, regions with low (AA 200-250) and high (AA 270–300) values of the pathogenicity score *REVEL* correspond to local minima and maxima of the curve. The heat-strip representation displays mean score values allowing a more fine-granular resolution (Fig. 2E).

#### 3.2.5. Statistical comparisons

Pathogenicity score values within the four variant groups *ClinVar _pathogenic, ClinVar_benign, gnomAD*, and *InSilico* are shown as violin plots with integrated quartiles (for definitions see Methods section 2.3). Additionally, score value distributions are statistically compared in order to assess the capability of the specific score to discriminate between variants of the different categories and hence its possible suitability for variant classification. For example, *ASS1* variants, that were annotated as P/LP, yield significantly higher *CADD* and *REVEL* score values than variants in the other three groups (Fig. 2F).

#### 3.2.6. Interactive table

On the bottom side of the user interface, an interactive table, that remains sticky during scrolling, is available (Fig. 1). It comprises two tabs with all ClinVar variants as well as all simulated nsSNVs and corresponding pathogenicity score values. In order to provide interactivity to the user, selected variants are displayed in the UTP and protein plot. Both tables can be filtered, e.g., by variant type. Individual cells with score values in the *in silico* table are color-coded according to the predicted variant effect using score-specific cut-offs.

## 4. Discussion

The availability of databases with clinical and genetic information has never been greater than it is today. Scientific and medical advances, particularly in terms of sequencing and storage capabilities, will lead to an exponential growth of information in the coming decades. However, database queries often require bioinformatic tools, which ultimately limit the yield and usability of such. To enable clinicians, scientists, and other users without prior bioinformatic knowledge to explore rich yet complex datasets, user-friendly tools with an intuitive interface and the possibility to easily export data for further processing are needed. Web server applications allow users to make such queries regardless of the device and operating system. aRgus is therefore designed as a lightweight, multidimensional R/Shiny application to enable fast database queries.

aRgus uses minimal user input in the form of the gene name according to HUGO Gene Nomenclature Committee (HGNC) standard. aRgus can thus retrieve information of variable complexity on the localization and distribution of pathogenic variants at the chromosomal and protein levels, which can be used to explore biological and biochemical properties, such as mutational hotspots of pathogenic and benign variance within proteins. Visual linkages of pathogenic variation can be generated by annotating functionally important regions and domains from the UniProt database. aRgus provides simple means of displaying complex distributional information using complexity-reduced density representation that is quick and easy for the human eye to comprehend. The user is offered a wide range of possibilities to select relevant information to answer respective research questions.

By allowing simultaneous display of variants stored in gnomAD, the issue of survivorship bias, as a form of selection bias, can be overcome. Survivorship bias occurs in all clinical genetic databases and potentially leads to oversight of variants, that did not pass biological selection, by sole assessment of pathogenic variants from clinical databases such as ClinVar. This often results in misconceptions in the interpretation of mutational hotspots. The gnomAD database v2.1 contains over 125,000 exomes and 15,000 genomes from different populations. A comparison of benign variants derived from gnomAD and pathogenic variants listed in ClinVar and other genetic databases thus enables an improved assessment of putative pathogenic hotspots on the gene and protein level.

Beyond pure visualization of information on known pathogenic variants, a polynomial regression model and heatmap visualization offer an additional way of data exploitation which can be particularly advantageous for proteins that have previously been described to only a limited extent. These models overcome inaccessible, tabular data on pathogenicity scores and simplify the comprehensibility of visualized predicted variant effects up to single amino acid resolution. By annotation of all biologically possible missense variants using 36 different pathogenicity scores, statements can be made about protein regions with high impact of amino acid exchanges without existing *in vitro* studies. Alternatively, resulting information can be used to plan functional *in vitro* studies, e.g., in order to investigate the functional relevance of regions in scarcely described proteins or with only limited data on pathogenic variants.

### 4.1. Limitations

Despite of its scientific value, aRgus is subject to some limitations. The quality of the visualizations and analyses produced by aRgus heavily depends on the quality of data available. According to our use cases, ClinVar data does not represent the entirety of all previously reported pathogenic variants. This is largely due to the lack of obligation of genetic laboratories to enter newly discovered disease-causing variants in centralized repositories. Extensive literature reviews are therefore necessary to obtain a comprehensive picture of mutational distribution. This could be significantly improved by the addition of further, commercial databases such as HGMD or LOVD (19, 20). To enable users to visualize variants identified through their own literature research or genetic studies, variants can be selected from the dbNSFP-derived table of pathogenicity score values and are automatically highlighted in all plots.

### 4.2. Conclusion

Combining accessible and interactive visualizations of genetic and variant data with pathogenicity analysis in a synoptic, standalone tool, aRgus outstands existing applications for genetic data exploitation regarding output versatility and flexibility (5, 21). With each update of the databases connected to aRgus, the diversity and analysis capabilities of its visualizations and datasets will also improve. Thus, aRgus will provide useful and previously mostly inaccessible information to a broad usership with limited bioinformatics skills such as practicing clinicians, basic scientists, and geneticists, and thus be helpful to answer scientific questions.

## 5. Acknowledgments

## 6. Author contributions

JS, TD, and HB devised the project and main conceptual ideas and designed the study. JS, HB, JH, AJ, SU, and DH have designed and delivered the technical realization and implementation of aRgus. All authors were involved in the further development of aRgus during the development period through their intellectual input and the execution of targeted analyses. All authors provided critical feedback and helped shape the research, analysis, and manuscript.

## 8. Supplementary files

### 8.1. Figure S1

**Fig. S1:**
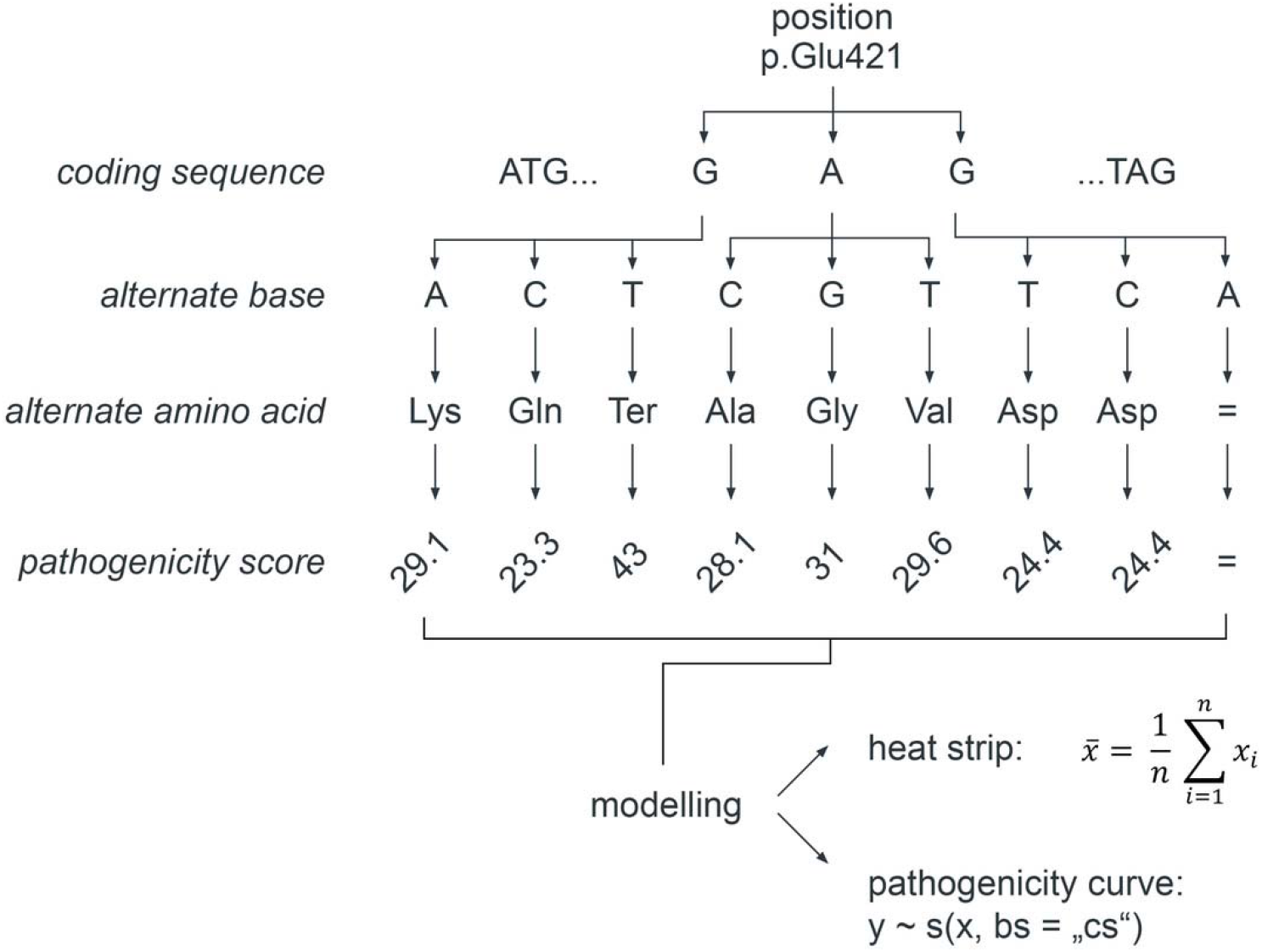
Schematic illustration of dbNSFP-derived variant simulation and aRgus-mediated visualization. Starting from the coding sequence of a gene transcript, any base at any position is exchanged with its three non-synonymous alternate bases (top). Individual pathogenicity score values (bottom) are assigned to the corresponding amino acid substitutions (middle). In aRgus, the resulting tabular data is modelled and visualized using a dual approach with a polynomial regression curve and a heat strip.

**8.2. Table S1.**
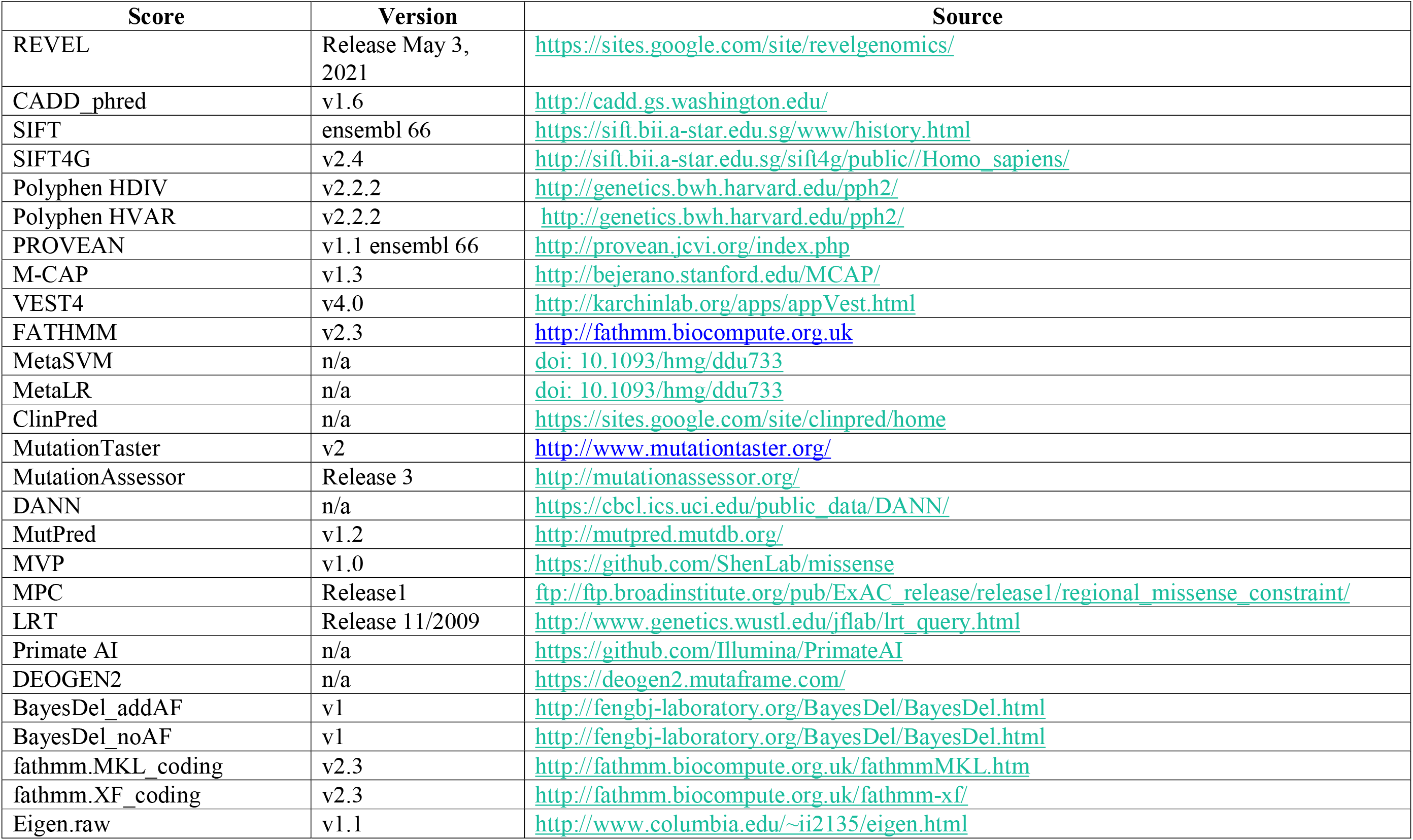
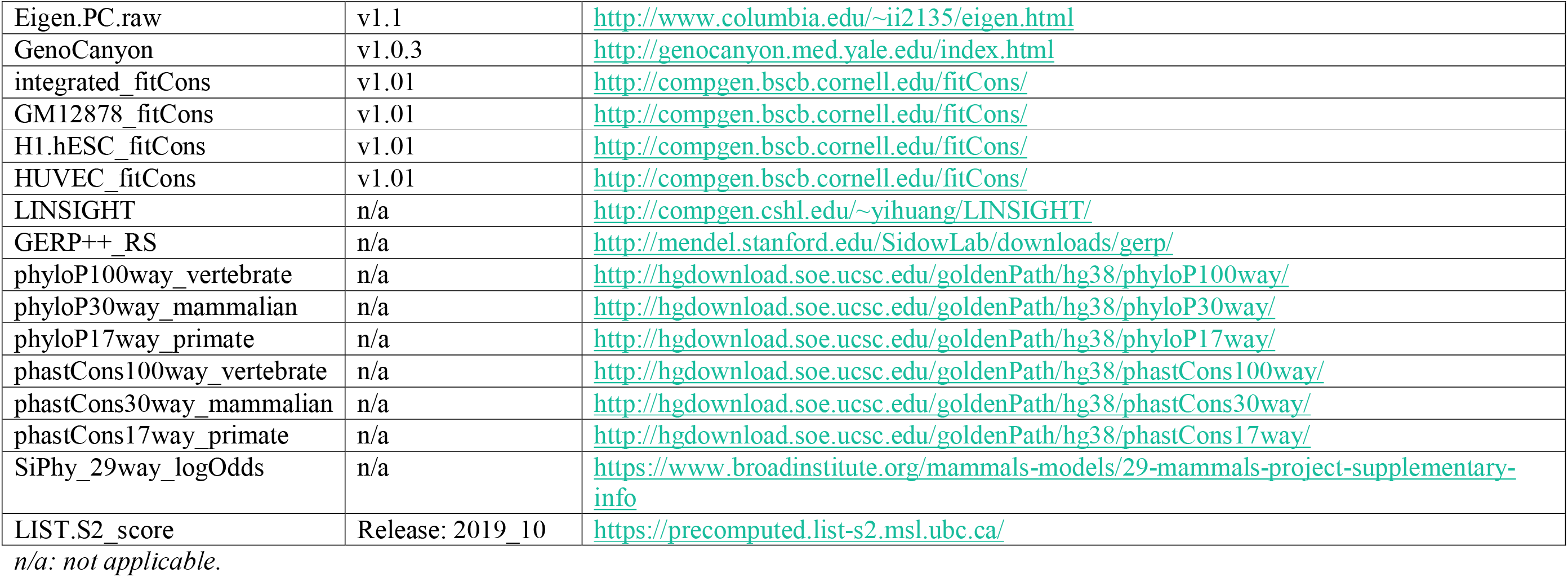
Pathogenicity scores available on aRgus.

